# Computational Prediction of Recombinant Expression Efficiency for Industrial Enzymes: An Integrated Co-Translational Sequence Analysis Platform

**DOI:** 10.64898/2026.10.07.757301

**Authors:** Tanushree Jaitly, Nicholas R. Silvaggi, Harry R. Horn, Raminderpal Singh, Edward Corrigan, John F. Conway

## Abstract

Industrial biocatalysis relies on the cost-effective production of active enzymes in heterologous hosts, yet a substantial fraction of candidate sequences fails during recombinant expression because sequence-level determinants of translational efficiency are not captured by conventional codon optimization alone. Here we present an integrated computational platform for pre-clinical assessment of industrial enzyme expression potential that combines baseline sequence metrics with co-translational modeling of ribosome stalling, translation rhythm, codon harmonization, folding bottlenecks, solubility proxies, and proteolysis risk. The platform accepts DNA, protein, FASTA, and PDB inputs and returns interpretable expression risk assessments, component-level diagnostics, variant design proposals, and actionable engineering recommendations. Analyses are parameterized for industrial expression hosts (Escherichia coli, Saccharomyces cerevisiae, Bacillus subtilis, and mammalian cells) and for native enzyme sources including Streptomyces, Bacillus, filamentous fungi, actinomycetes, and extremophiles. All analyses execute locally without dependence on external databases, enabling rapid and secure screening of proprietary enzyme candidates prior to gene synthesis and fermenter scale-up. We report three layers of preliminary validation on small cohorts: (i) a retrospective public industrial-enzyme ranking benchmark (n = 20; rank subset n = 10); (ii) calibration of the solubility module on held-out eSOL enzymes with quantitative experimental solubility; and (iii) an independent collaborator wet-lab construct set retained as a true external-test holdout and never used for model fitting (n = 38 labeled constructs). On the external holdout, solubility calls achieved accuracy 68.4% (95% CI 52.5–80.9%), sensitivity 56.5% (36.8–74.4%), specificity 86.7% (62.1–96.3%), balanced accuracy 71.6% (bootstrap 95% CI 57.4–84.4%), and MCC 0.43 (0.15–0.69). Sensitivity remains close to chance on this small set, whereas specificity is higher; intervals are wide and the evidence should be read as directional rather than definitive. A directional arginine-surface redesign example (SrMilN insoluble parent to SrMilN_SASAR partially soluble variant; incomplete rescue with multiple soluble bands) was recovered. These findings support preliminary screening utility and transparent experimental agreement, but do not constitute high-accuracy prediction of industrial fermenter titre. Offline EC-class heuristics are retained only as optional provisional labels and are not presented as a validated platform capability (agreement 3/20 = 15%, 95% CI 5.2–36%; chance for six classes ≈ 16.7%).

## Introduction

Enzymes are central to industrial biotechnology, enabling biocatalytic processes in pharmaceuticals, food and beverage manufacturing, detergents, biofuels, and fine chemicals with reduced energy demand and improved stereoselectivity relative to many chemical routes (Schmid et al., 2001; Bornscheuer et al., 2012). The commercial viability of an industrial enzyme program depends not only on catalytic performance under process conditions but also on the ability to produce the protein at sufficient titre, solubility, and stability in a scalable recombinant host. Despite decades of progress in molecular cloning and fermentation engineering, heterologous expression remains a primary bottleneck: sequences that are abundant and functional in their native producer particularly actinomycetes, streptomycetes, and filamentous fungi frequently express poorly when transferred to standard industrial platforms such as Escherichia coli or yeast (Anne et al., 2014; Song et al., 2017).

Traditional approaches to improving expression emphasize synonymous codon replacement to maximize codon adaptation index (CAI) or to eliminate rare codons in the production host (Sharp and Li, 1987; Gustafsson et al., 2004). While CAI optimization can increase yield for some targets, industrial enzyme projects increasingly recognize that translation is a dynamic, co-translational process in which elongation rate heterogeneity influences nascent-chain folding, aggregation propensity, and downstream proteolytic susceptibility (Zhang et al., 2009; Pechmann et al., 2013; Willmund et al., 2013). Arginine-rich segments and consecutive arginine dipeptides (RR motifs), for example, have been associated with ribosome pausing and reduced soluble recovery of recombinant proteins, including enzymes bearing positively charged surface patches derived from natural biodiversity (Tuller et al., 2010; Chu et al., 2014). Similarly, abrupt transitions in codon rarity along a coding sequence rather than average adaptation alone can create local slowdowns that promote misfolding of large, multi-domain industrial enzymes during synthesis (Cannarozzi et al., 2015).

Industrial enzyme discovery pipelines now generate large numbers of candidate sequences from metagenomes, genome mining, and protein engineering campaigns. Experimental expression testing of every variant across multiple hosts is costly and slow. There is therefore a need for computational tools that translate sequence information into expression risk assessments aligned with bioprocess decision-making: which constructs to synthesize first, which host to prioritize, and which sequence modifications are most likely to improve manufacturability before commitment to fermenter scale-up (Kumar et al., 2018). Existing web servers and commercial gene-design tools often treat expression as a single scalar optimization problem and may not surface co-translational failure modes relevant to enzyme classes with complex cofactor requirements, disulfide connectivity, or high GC content typical of bacterial secondary metabolite producers.

To address this gap, we developed a modular computational platform for co-translational enzyme expression engineering that evaluates industrial enzyme sequences through a unified analytical pipeline. The platform integrates host-aware codon usage analysis, GC content classification, RR and poly-arginine motif detection, organism signature inference, surface arginine engineering, codon harmonization, translation rhythm modeling, ribosome stall prediction, co-translational folding bottleneck identification, solubility and aggregation patch analysis, proteolysis risk scoring, variant design proposals, and composite expression scoring with explicit recommendations. An offline EC-class motif heuristic is available as an optional provisional label for workflow convenience, but it is not marketed as a predictive capability in this preprint because current panel agreement is at chance (see Results). The system is designed for enzymes intended for industrial biocatalysis and operates entirely on local sequence analysis without mandatory calls to paid annotation services, supporting secure evaluation of proprietary sequences.

In this work, we describe the scientific rationale and conceptual analytical workflow of the platform with emphasis on industrial enzyme applications. We demonstrate representative output on the authentic Streptomyces hygroscopicus α-amylase coding sequence (UniProt P08486; GenBank M15540) targeted for heterologous production in E. coli, report retrospective ranking of public industrial enzymes, calibrate solubility against experimental eSOL measurements, and evaluate an independent collaborator wet-lab holdout. Proprietary scoring parameters and implementation details are intentionally withheld.

## Methods

### Platform overview and scope

The platform is a modular sequence-analysis system that assesses recombinant expression risk for industrial enzyme candidates prior to gene synthesis and scale-up. Users submit enzyme sequences through a secure interface and receive structured, interpretable reports. Supported input modalities include nucleic acid and protein sequences, standard FASTA records, and optional structural coordinate files from which polypeptide sequence can be derived (Berman et al., 2000). Input type may be specified by the user or inferred automatically from sequence composition. When coding DNA is provided, translation to protein is performed using the standard genetic code for motif and classification analyses.

Analyses are contextualized by expression host and, optionally, by native source organism. Supported industrial hosts include Escherichia coli, Saccharomyces cerevisiae, Bacillus subtilis, and mammalian systems, reflecting dominant production platforms in industrial biotechnology (Rosano and Ceccarelli, 2014; Baeshen et al., 2015; Demain and Vaishnav, 2009). Source organism context (for example Streptomyces, Bacillus, fungal, actinomycete, or extremophile backgrounds) informs codon-harmonization and rhythm analyses when relocating genes from native producers to heterologous factories (Anne et al., 2014; Ferrer et al., 2016). Host- and source-specific reference profiles are maintained internally and are not disclosed here.

### Analytical framework

Each submitted sequence is processed through a unified multi-module pipeline that combines baseline sequence descriptors with co-translational manufacturability models (Kumar et al., 2018; Hidalgo et al., 2022; Netzer and Hartl, 2016; Zhang et al., 2009). Modules operate on the same input and contribute to complementary summary scores and engineering recommendations. Exact algorithmic formulations, decision thresholds, and composite weightings constitute proprietary intellectual property and are omitted from this preprint.

### Baseline sequence descriptors

Arginine-rich motif analysis: Protein sequences are examined for consecutive arginine dipeptides and extended poly-arginine stretches associated in the literature with ribosome pausing and reduced soluble recovery (Tuller et al., 2010; Chu et al., 2014; O’Brien et al., 2014). Results are summarized as categorical risk levels and positional annotations to guide construct redesign.

Host-aware codon usage: For DNA inputs, in-frame codon composition is compared against host-specific rare and preferred codon references derived from published codon-usage principles and industrial expression practice (Sharp and Li, 1987; Welch et al., 2009; Gustafsson et al., 2004; Plotkin and Kudla, 2011). Outputs include adaptation-oriented summary metrics and rare-codon burden indicators. Protein-only submissions omit codon analysis and rely on other modules.

GC composition: Overall guanine-cytosine content is computed and classified relative to composition ranges relevant to cloning and heterologous expression, particularly for GC-rich genes from actinomycete and fungal enzyme sources (Bernardi, 2000). Extreme composition contributes to manufacturability risk assessment.

Organism signature inference: When DNA is available, compositional and codon-distribution features are compared to reference organism signatures to estimate whether the sequence resembles the selected expression host or a phylogenetically distant source (Sharp and Li, 1987; Karlin et al., 2001; Bentley et al., 2002). Host-source mismatch triggers harmonization-oriented recommendations.

Optional provisional EC-class labels (not a claimed capability): Sequences can be assigned provisional Enzyme Commission class labels (classes 1–6) using internal motif and composition heuristics for offline workflow labeling without mandatory external homology searches (Bairoch, 2000; Webb et al., 2022). In the present industrial panel this heuristic performs at chance (≈15% first-level agreement versus ≈16.7% expected for random six-class guessing) and must not be treated as EC annotation, regulatory support, or a manufacturability feature. Users should supply curated UniProt, BRENDA, or InterPro EC numbers for any reporting use (UniProt Consortium, 2023). Specific motif libraries and scoring rules are proprietary and are not listed here.

### Co-translational and manufacturability modules

The following modules extend baseline descriptors to capture translational and folding constraints relevant to industrial enzyme production. Implementation details, parameterizations, and scoring equations are withheld.

Structural context: Sequence-based domain and secondary-structure propensity estimates, optionally informed by supplied structural coordinates, provide context for folding and solubility modules (Kyte and Doolittle, 1982; Chou and Fasman, 1978; Berman et al., 2000).

Surface arginine engineering: Positively charged surface features are identified and conservative substitution options are proposed to mitigate polybasic pausing and aggregation risk while preserving functional constraints where possible (Davis et al., 1999; Trevino et al., 2007).

Codon harmonization: Rather than maximizing average codon adaptation alone, harmonization evaluates local rare-codon patterning and pause-site utility, recommending synonym replacements that improve host compatibility while preserving beneficial translational rhythm (Cannarozzi et al., 2015; Kudla et al., 2009).

Translation rhythm modeling: Local codon-rarity profiles are used to highlight abrupt deceleration regions that may precede folding bottlenecks in large industrial hydrolases and oxidoreductases (Tuller et al., 2010; O’Brien et al., 2014).

Ribosome stall prediction: Polybasic runs, arginine-rich motifs, and local rarity spikes are integrated to estimate stall propensity along the nascent chain (Chu et al., 2014; Tuller et al., 2010).

Co-translational folding analysis: Regions where slow translation may coincide with domain emergence are flagged as elevated aggregation risk during synthesis (Willmund et al., 2013; Pechmann et al., 2013; Netzer and Hartl, 2016).

Solubility and aggregation proxies: Hydrophobicity, charge, merged hydrophobic-patch coverage, arginine composition, and related sequence descriptors are used as proxies for inclusion-body risk (Kyte and Doolittle, 1982; Hidalgo et al., 2022; Magnan et al., 2009). In the current implementation, these features feed a calibrated linear solubility model trained on reviewed eSOL enzymes with quantitative chaperone-free solubility measurements; DeepSol records were retained only as secondary evaluation and were not used for fitting. Collaborator wet-lab constructs were reserved exclusively as an external-test holdout.

Proteolysis and truncation risk: Flexible or protease-susceptible regions associated with truncated or low-titre products are annotated (Guruprasad et al., 1990; Terpe, 2006).

Variant design: The platform proposes a small set of prioritized redesign strategies (for example surface-charge edits and rhythm-preserving recoding concepts), with qualitative rationales to support gene-synthesis ordering. Synthesis-ready full recoded DNA output may be restricted depending on deployment configuration.

### Composite scoring and reporting

Module outputs are summarized as complementary composite expression-risk scores and qualitative performance bands to support construct prioritization. Scores are presented with component-level diagnostics so that users can identify which biological factors most limit predicted manufacturability. Recommendation text is generated when modules detect elevated arginine-related risk, codon-host mismatch, unfavorable composition, host–source discordance, folding or solubility concerns, or related manufacturability flags. Interactive visualizations (for example rarity profiles, component summaries, and stall maps) are available in the user interface. Exact band boundaries, component normalizations, and weighting schemes are proprietary and are not published.

### Retrospective validation design

Validation was organized in three layers. First, a retrospective public industrial-enzyme ranking benchmark tested whether the integrated co-translational score ordered enzymes consistently with published heterologous-expression difficulty; heuristic first-level EC assignment was recorded only as a negative control / non-capability check against curated annotation, not as a primary performance claim. The ranking cohort comprised 20 unique industrial or biotechnology enzymes from bacterial, fungal, extremophile, and mammalian sources. Case-study views in the application were presentation layers derived from this cohort and were not counted as additional validation records. Second, the solubility module was calibrated on reviewed eSOL enzymes with measured solubility and evaluated on a held-out eSOL split. Third, an independent collaborator wet-lab construct set with binary Soluble/Insoluble outcomes was evaluated as a true external_test holdout and was never used to fit solubility or integrated-score weights. Because ranking (n = 20 / rank-subset n = 10) and external (n = 38) cohorts are small and non-independent of project history, all point estimates are reported with 95% confidence intervals.

For the public ranking cohort, protein sequences and source-organism metadata were retrieved from the UniProt REST API (UniProt Consortium, 2023). Expected EC numbers were cross-referenced with UniProt and BRENDA (Chang et al., 2021), and CAZy annotations were used as supporting metadata for glycoside hydrolases (Drula et al., 2022). Published expression reports were used to assign qualitative heterologous-expression tractability categories in an E. coli context (tractable, condition-dependent, difficult). eSOL provided quantitative solubility supervision for model calibration (Niwa et al., 2009; Niwa et al., 2012). The collaborator external set comprised project constructs with experimental soluble/insoluble labels, including arginine-surface redesign variants.

Each public ranking sequence was analyzed through one unified workflow comprising baseline sequence analysis followed by integrated co-translational analysis. Predicted first-level EC class was compared with the curated class solely to quantify heuristic failure relative to chance. For a prespecified 10-record rank subset, tractability categories were encoded as tractable = 3, condition-dependent = 2, and difficult = 1. Ranking endpoints were category-mean integrated scores, the directional contrast between tractable and difficult groups, and Spearman correlation with an approximate Fisher-z 95% interval (acknowledging instability at n = 10). For calibrated solubility, held-out eSOL performance was summarized by Spearman/Pearson correlation and balanced accuracy at a 50% solubility threshold. For the collaborator external test, we report accuracy, sensitivity, specificity, precision, balanced accuracy, Matthews correlation coefficient (MCC), and calibration by predicted-score bin, each with 95% Wilson intervals for binomial rates and bootstrap percentile intervals (10,000 resamples) for MCC and balanced accuracy. Unlabeled collaborator variants were excluded from metric calculation.

### Definition of expression-tractability categories

Tractable records were those for which the literature described substantial soluble or active recombinant protein under standard expression conditions with little sequence or process intervention. Condition-dependent records were those for which expression was achievable but depended on measures such as reduced induction temperature, codon adaptation, fusion tags, chaperone support, cofactor supplementation, secretion engineering, or partial refolding. Difficult records were those associated with predominantly insoluble or inactive product, very low soluble recovery, extensive refolding, complex disulfide or glycosylation requirements, or major source–host incompatibility. These categories were qualitative because the source publications did not report yield using a common assay or unit; they must not be interpreted as fixed concentration thresholds.

Two records were selected for detailed illustration: Bacillus licheniformis α-amylase as a familiar industrial hydrolase from the tractable category and Taq DNA polymerase as a thermophilic enzyme from the condition-dependent category. They were selected to provide contrasting source biology and module outputs, not as independent evidence of accuracy. Category names are disclosed here to make selection transparent but are omitted from the case-study titles.

### Software and confidentiality

The platform is delivered as a secure, locally executable analytical service with a browser-based report interface. Host codon profiles, motif libraries, classification heuristics, composite weights, and related configuration assets are proprietary and are not redistributed with this preprint. No external paid annotation services are required for core analyses. Machine-learning models were not required for the analyses reported here, although ML-based expression predictors remain an active area of research (Hidalgo et al., 2022; Cao et al., 2021).

### Interpretation guidelines and limitations

All predictions are intended for pre-synthesis decision support in industrial enzyme programs and do not replace experimental expression testing, commercial gene design, or authoritative EC annotation (Rosano and Ceccarelli, 2014; Demain and Vaishnav, 2009). Organism predictions and any optional EC-class labels should be confirmed with curated databases; EC heuristics are explicitly not a validated capability of this platform. Users should prioritize remediation of the weakest diagnostic categories and validate top-ranked variant proposals in micro-scale expression before fermenter scale-up (Terpe, 2006). Dataset limits should be stated explicitly in partner communications: primary quantitative calibration used eSOL cell-free E. coli measurements rather than intact-cell industrial fermentation; most public training sequences lack coding DNA, so codon/rhythm modules remain weakly supervised; the collaborator external set is small (n = 38) with wide confidence intervals and project enrichment; sensitivity is only modestly above chance on that set; binary soluble/insoluble labels omit quantitative yield and activity; and EC-class prediction remains at chance and informational only.

### Data availability

The validation panel, record-level results, aggregate summary, and case-study outputs are included with the project as validation/panel/starter_panel.csv and validation/results/. External wet-lab holdout metrics and the SrMilN directional-example summary are provided in validation/wetlab/EXTERNAL_WETLAB_REPORT.md and validation/wetlab/external_test_metrics.json. Protein sequences and accession metadata for public records are retrievable from UniProt. BRENDA was used for EC cross-reference and CAZy for supporting glycoside hydrolase metadata; neither database was redistributed. Collaborator sequences and labels are confidential and are not redistributed. Proprietary scoring parameters, reference profiles, motif libraries, and source code are not publicly available. Result figures corresponding to Figures 1 and 2 in the manuscript are provided as validation/figures/figure1_experimental_agreement.png and validation/figures/figure2_external_highlights.png (PDF versions also available); numerical summaries corresponding to Tables 1–3 are reproduced in the Results section.

**Table 1.** Retrospective benchmark of the unified expression-analysis workflow.

| Endpoint | Result |
| --- | --- |
| Unique records analyzed successfully | 20/20 (100%) |
| Rank subset | 10 records: 2 tractable, 5 condition-dependent, 3 difficult |
| Mean integrated score: tractable | 47.17 |
| Mean integrated score: condition-dependent | 37.01 |
| Mean integrated score: difficult | 32.22 |
| Directional contrast | Tractable > difficult (pass) |
| Spearman correlation | $\rho = 0.458$ (approx. 95% CI $-0.24$ to $0.84$; $n = 10$) |
| EC-class agreement (negative control; not a capability) | 3/20 (15%; 95% CI 5.2–36%; chance $\approx 16.7\%$) |

**Figure 1.**
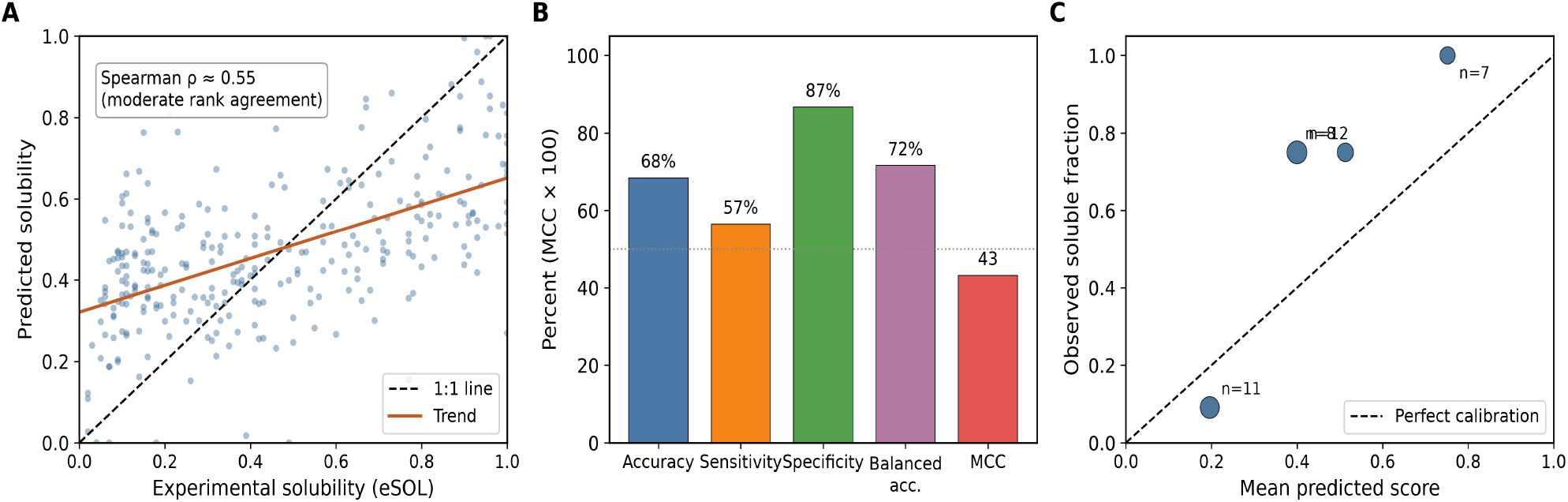
Experimental agreement of the calibrated solubility model (plain-language guide). (A) Each point is one held-out eSOL enzyme (n = 294). The x-axis is experimental solubility; the y-axis is the model prediction. The dashed line is a perfect match; the solid line is the overall trend. The scatter looks broad by eye, but Spearman’s rank correlation (ρ ≈ 0.55) shows moderate ranking agreement: enzymes that are more soluble experimentally tend to receive higher predicted scores (ρ = 0 would mean no ranking signal; ρ = 1 would mean perfect order). This supports triage, not fermenter-titre prediction. (B) External collaborator wet-lab holdout (n = 38). Accuracy = percent of all soluble/insoluble calls that matched the lab; sensitivity = percent of truly soluble constructs correctly flagged soluble; specificity = percent of truly insoluble constructs correctly flagged insoluble; balanced accuracy = average of sensitivity and specificity; MCC = chance-corrected agreement. (C) Calibration: constructs were grouped by predicted score; the plot asks whether higher score bins contain a higher fraction of lab-soluble proteins (marker size reflects bin count).

**Figure 2.**
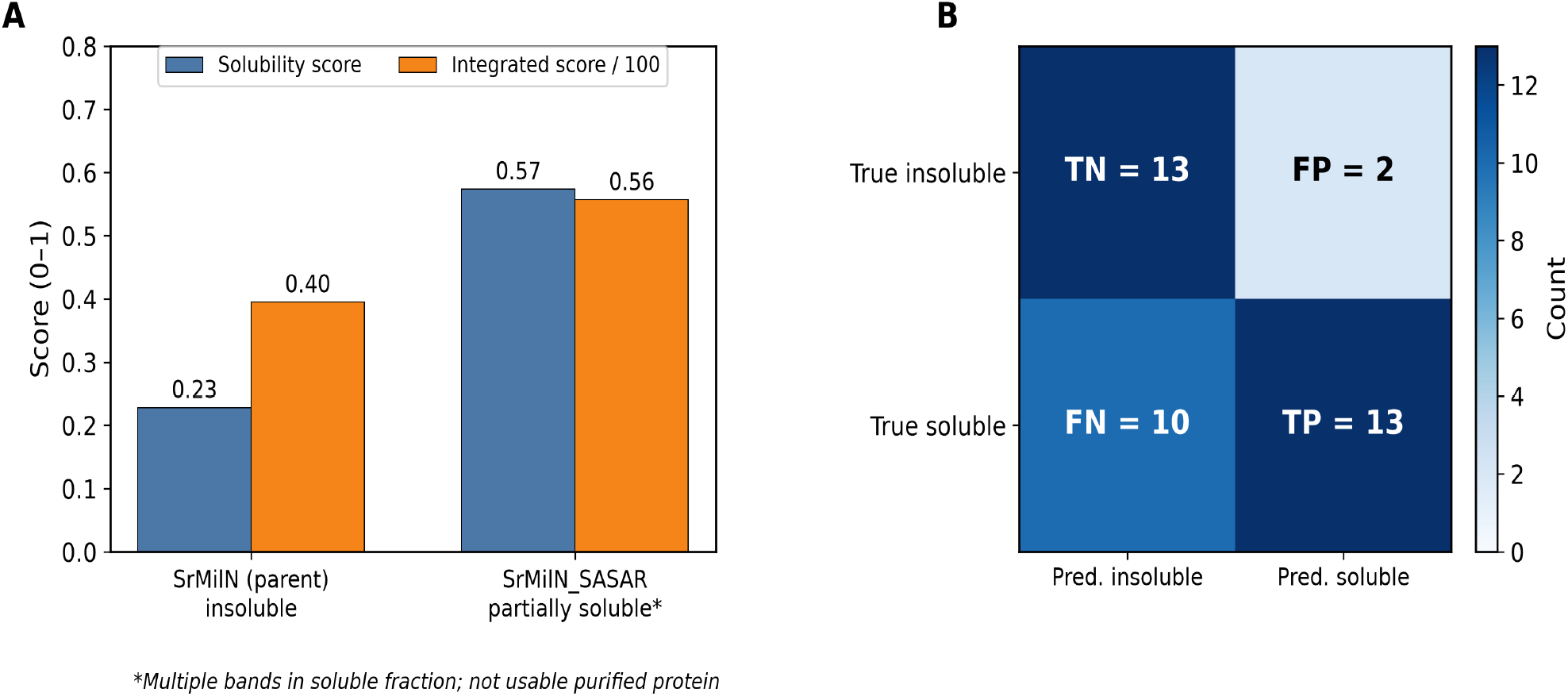
External wet-lab validation highlights (plain-language guide). (A) Directional score shift for an arginine-surface redesign: parental SrMilN was insoluble in the wet lab; SrMilN_SASAR became partially soluble (a family of bands in the soluble fraction) but was not purification-ready or usable as clean single-species protein. Both the solubility score and the integrated expression score increased for the redesigned variant - a directional improvement, not a process-ready success. (B) Confusion matrix for the external holdout (n = 38; threshold 0.46). TP = correctly called soluble; TN = correctly called insoluble; FP = false soluble alarm; FN = missed soluble construct.

## Results

### Representative analysis of Streptomyces hygroscopicus α-amylase (UniProt P08486)

To illustrate platform output on real biology, we analyzed the complete coding sequence of Streptomyces hygroscopicus α-amylase (EC 3.2.1.1; UniProt P08486; GenBank/EMBL M15540, CDS coordinates 264–1700), a published actinomycete industrial hydrolase frequently discussed in the context of difficult heterologous expression in E. coli (Hoshiko et al., 1987). The submitted input was the authentic 1,437-nt CDS (479 codons including the terminal stop), which translates in-frame to the 478-residue UniProt mature/precursor protein sequence. Source context was set to Streptomyces and the expression host to E. coli. The platform returned a unified report spanning baseline sequence metrics, co-translational manufacturability assessment, variant design proposals, and engineering recommendations within a single analysis run.

### Input sequence

The complete CDS is publicly available under GenBank M15540 and UniProt P08486; reproducing the full 1,437 nt in the manuscript is unnecessary. The 5′ and 3′ termini of the analyzed CDS are shown below to document the exact construct boundaries used. >P08486_M15540_Streptomyces_hygroscopicus_alpha_amylase (CDS 1,437 nt; aa 1–478) 5′-ATGCAGCAACGTTCCCGTGTGCTGGGCGGGACGCTCGCCGGAATAGTGGCCGCGGCGGCGGCCACC……GCGCTGCACGTCCCCGGTCAGTCGGCCGGGTCGCCCCGCTCCAGCGCCAAGAGGGTGGAGCAGTAG-3′Translated N-terminus: MQQRSRVLGGTLAGIVAAAAATVAPWPSQATPPGQKTVTATLFER… (478 aa; matches UniProt P08486).

### Sequence parsing and identity

The sequence was detected as FASTA DNA and translated for motif and EC analyses, yielding the full 478-amino-acid product that matches the UniProt P08486 sequence exactly. GC content was 70.01% (1,006 G/C, 431 A/T), classified as high and typical of Streptomyces coding sequences. Four hundred seventy-nine complete codons were parsed (478 sense codons plus the terminal stop). The sequence identifier was retained from the FASTA header and linked to the public accessions above.

### Baseline sequence metrics

Arginine-motif analysis returned no consecutive RR dipeptides and no poly-arginine regions (risk: low), indicating no immediate arginine-mediated stalling burden at the dipeptide level. Codon usage relative to the E. coli profile identified a rare-codon fraction of 3.13% (15 of 479 codons) with a low host-adaptation (CAI-like) score of 0.273, consistent with GC-rich actinomycete codon bias poorly matched to E. coli. Organism-signature inference predicted E. coli-like compositional features with only moderate confidence (66.3%) driven by codon-usage signals under the forced host context. An optional offline EC-class heuristic incorrectly assigned Class 5 (Isomerases) rather than the curated hydrolase annotation (EC 3.2.1.1), consistent with the at-chance panel performance of this label; curated UniProt/BRENDA EC numbers should be used for any reporting.

### Co-translational and manufacturability metrics

Ribosome stall prediction indicated low stall risk, with no enumerated stall sites and no polybasic runs. Solubility analysis indicated a fair solubility band (score 0.37; GRAVY −0.30; net charge at pH 7 approximately −4.9), with 14 merged hydrophobic aggregation patches distributed along the nascent chain, including an N-terminal hydrophobic segment spanning residues 7–25. Codon harmonization returned a high harmonization score (0.93) under a rhythm-preserving strategy, flagged 13 pathological rare codons (predominantly CCC, with CGA and AGG contributions) for replacement with E. coli-preferred synonyms, and preserved 2 beneficial pause sites at putative domain boundaries. Translation rhythm modeling indicated relatively coherent velocity structure along the CDS (rhythm score 0.85). Co-translational folding analysis indicated substantial bottleneck burden (folding score 0.14; 20 hydrophobic-cluster bottlenecks), consistent with early hydrophobic segments that may aggregate if translation is too fast. Surface arginine analysis identified 21 solvent-exposed arginines in 2 clustered patches (engineering priority: moderate) and recommended conservative R→K/Q/S/T substitutions at exposed positions. Proteolysis/truncation risk was high.

### Composite scores and engineering recommendations

The baseline expression score was 59.1 (band: fair), driven primarily by favorable arginine-motif status offset by weak codon adaptation to E. coli and high GC content outside the preferred 40–60% window. Baseline recommendations included tuning GC toward a more balanced composition and reviewing class-specific expression constraints - recommendations that must be interpreted against the curated EC 3.2.1.1 hydrolase annotation rather than the provisional isomerase call.

The integrated co-translational expression score was 49.8 (band: good on the calibrated v2 threshold scale; not directly interchangeable with baseline bands). Primary limiting factors were co-translational folding (component 0.14), solubility (0.37), and proteolysis risk, despite favorable stall (1.0), harmonization (0.93), and rhythm (0.85) components. Additional recommendations included surface arginine engineering at exposed clusters, reduction of hydrophobic aggregation patches (for example N- or C-terminal fusion strategies), proteolysis/truncation checks by SDS-PAGE and N-terminal sequencing, and rhythm-aware rather than global-CAI codon redesign. The divergence between baseline (59.1) and integrated (49.8) scores highlights manufacturability constraints - folding bottlenecks and solubility patches - not captured by codon adaptation alone.

### Variant design proposals

The variant design module proposed a prioritized surface-arginine construct within a combined surface-arginine and rhythm-preserving harmonization design strategy:

- Variant 1 (surface arginine strategy; composite variant score 0.65): R4K, R6K, and R45K substitutions at solvent-exposed positions to reduce polybasic/surface aggregation risk, with DNA-level recoding required separately.
- Harmonization arm (rhythm-preserving strategy): replace the 13 pathological rare codons with E. coli-preferred synonyms while preserving the 2 beneficial pause sites; do not maximize CAI globally. This arm is recommended in combination with Variant 1 for gene synthesis planning.

Variant 1 was returned as the scored protein-level redesign, with combined surface arginine engineering and rhythm-preserving harmonization recommended before ordering a synthesis construct for micro-scale expression.

### Visualization outputs

The analysis returned interactive visualizations including a codon-rarity profile along the 1,437-nt coding sequence, a component summary comparing co-translational module outcomes, and a ribosome stall map indicating low stall risk under the current sequence context.

### Interpretation

This representative case illustrates a common industrial enzyme scenario: an authentic, full-length actinomycete α-amylase CDS with acceptable-to-moderate baseline metrics that nonetheless exhibits co-translational manufacturability constraints - high GC/codon mismatch, distributed hydrophobic aggregation patches, folding bottlenecks, and surface-arginine engineering opportunities - detectable through integrated analysis. For Streptomyces-to-E. coli relocation projects, CAI maximization alone would miss these failure modes. Prior to scale-up, we recommend synthesizing a harmonized CDS incorporating the R4K/R6K/R45K surface edits (or experimentally guided alternatives) and validating soluble titre in a micro-scale expression screen before fermenter commitment. Full analysis artifacts for this accession are archived with the project validation outputs.

### Retrospective benchmark of public industrial-enzyme records

All 20 unique panel records were retrieved from UniProt and analyzed successfully (100% processing success). The rank-validation subset contained two tractable, five condition-dependent, and three difficult records. Mean integrated scores decreased in the expected direction: 47.17 for tractable, 37.01 for condition-dependent, and 32.22 for difficult records. The prespecified directional contrast therefore passed (tractable > difficult). Spearman correlation between integrated score and the ordinal literature prior was ρ = 0.458 (approximate Fisher-z 95% CI −0.24 to 0.84; n = 10), indicating a moderate point estimate whose interval includes zero and therefore remains compatible with no rank association in this small retrospective subset. First-level EC-class agreement was 3/20 (15%, Wilson 95% CI 5.2–36%), indistinguishable from six-class chance (≈16.7%); accordingly we do not claim EC assignment as a platform capability. These ranking endpoints are summarized in Table 1.

### Calibrated solubility model on experimental eSOL enzymes

To address the lack of discrimination in the earlier solubility heuristic, a calibrated solubility model was trained on reviewed eSOL enzymes with quantitative chaperone-free solubility and evaluated on a held-out eSOL split (n = 294). DeepSol sequences were retained only as secondary evaluation and were not used for fitting. Held-out performance improved substantially relative to the previous heuristic (Spearman correlation approximately 0.55 versus 0.09; balanced accuracy approximately 0.73 versus 0.50 at a 50% solubility threshold). Figure 1A plots predicted versus experimental solubility for these held-out enzymes. To a non-specialist eye the scatter can look almost uncorrelated because points form a broad cloud rather than a narrow diagonal; that appearance is expected for biological solubility data. The relevant summary statistic is Spearman’s rank correlation (ρ ≈ 0.55): if enzyme A is more soluble than enzyme B experimentally, the model tends to assign A a higher score than B about as often as a moderate rank association would imply. Spearman ρ = 0 would be no ranking signal, ρ = 1 would be perfect rank order, and ρ ≈ 0.55 is moderate agreement - useful for triage, not a claim of tight 1:1 titre prediction. A dashed 1:1 line and a best-fit trend are shown in Figure 1A to guide the eye. These results support preliminary quantitative agreement with experimental solubility while remaining insufficient for manufacture-ready titre prediction.

### Wet-lab holdout

An independent collaborator construct set with experimental Soluble/Insoluble labels was evaluated as a true external-test holdout and was never used to fit model weights. After excluding unlabeled variants, 38 constructs remained. Using the eSOL-derived decision threshold (0.46), the platform made a binary “likely soluble / likely insoluble” call for each construct. This means in wet-lab setting: accuracy is the overall fraction of constructs whose soluble/insoluble call matched the experiment (68.4%; Wilson 95% CI 52.5–80.9%); sensitivity (true-positive rate) is the fraction of truly soluble constructs that the model correctly flagged as soluble (56.5%; 36.8–74.4%) - here only modestly above a coin flip; specificity (true-negative rate) is the fraction of truly insoluble constructs correctly flagged as insoluble (86.7%; 62.1–96.3%), the stronger signal on this set; precision is the fraction of “soluble” calls that were truly soluble (86.7%; 62.1–96.3%); balanced accuracy averages sensitivity and specificity so that the larger soluble class does not dominate (71.6%; bootstrap 95% CI 57.4–84.4%); and MCC (Matthews correlation, here 0.43; 0.15–0.69) is a chance-corrected agreement score that uses all four confusion-matrix cells (TP = 13, FP = 2, TN = 13, FN = 10), with 0 ≈ random and 1 = perfect. Table 2 consolidates these holdout metrics with intervals, and Figure 1B displays them graphically. Calibration was directional (Figure 1C): predicted scores below 0.30 were mostly insoluble (observed soluble fraction 0.09), whereas scores above 0.60 were soluble in this set (observed fraction 1.00). Mid-range scores were the principal source of false negatives, which is why Figure 1C matters for deciding when a high-specificity call is trustworthy versus when a mid-range score should trigger experimental follow-up rather than an automatic reject. On this small non-independent set the confidence intervals remain wide, so the evidence should be read as directional screening support rather than definitive predictive performance.

**Table 2.**
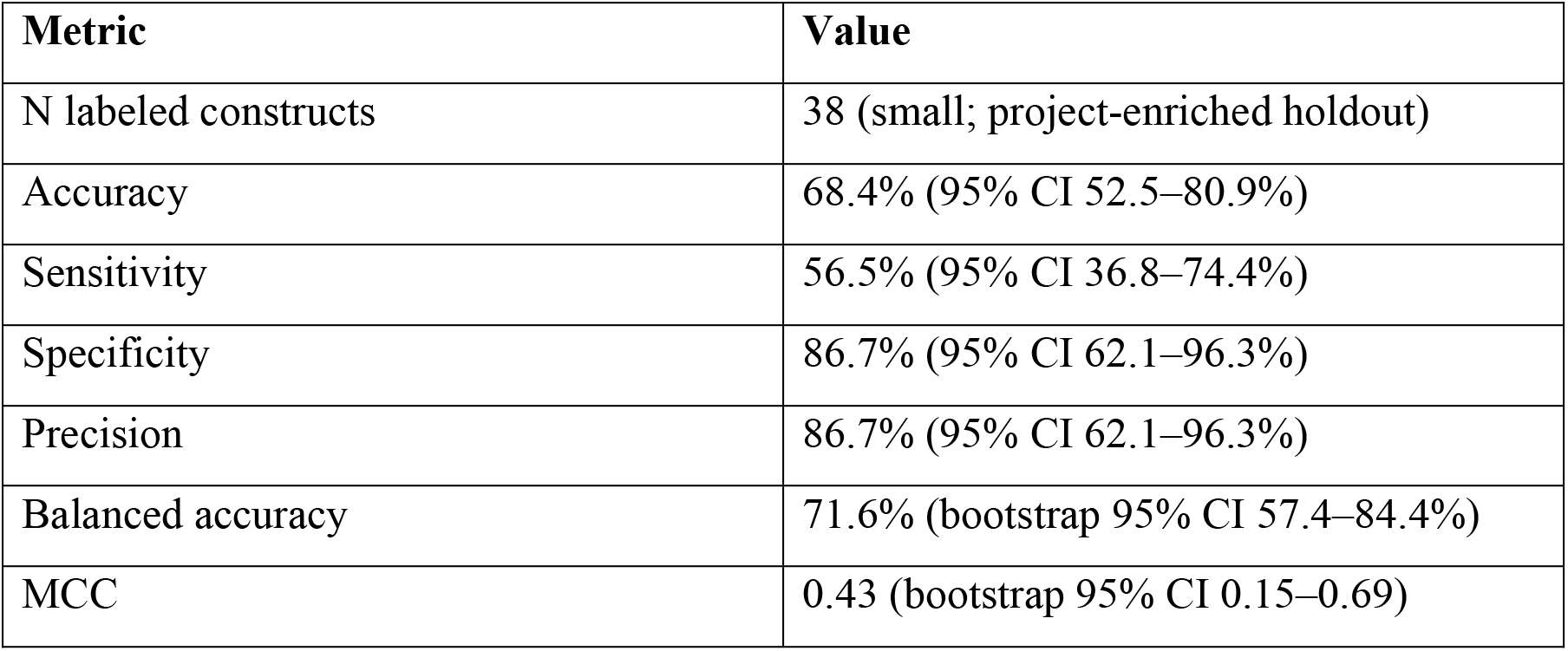
External wet-lab holdout performance of the calibrated solubility call (n = 38; 95% CIs).

Beyond aggregate metrics, partner evaluation also requires a concrete directional example - with carefully limited claims. Figure 2 and Table 3 therefore present a paired arginine-surface redesign from the same external holdout: parental SrMilN versus the engineered SrMilN_SASAR variant, together with the corresponding confusion matrix for the solubility call.

**Table 3.** Directional SrMilN arginine-surface redesign (partial solubility rescue; not purification-ready).

| Construct | Wet-lab outcome | Solubility score | Integrated score |
| --- | --- | --- | --- |
| SrMilN (parent) | Insoluble | 0.23 | 39.5 |
| SrMilN_SASAR | Soluble (not 100%) | 0.57 | 55.7 |

### Directional wet-lab success example: SrMilN arginine-surface redesign

A paired arginine-surface redesign provides a useful directional example rather than a claim of process-ready protein (Table 3; Figure 2A). The parental construct SrMilN was insoluble in wet-lab testing and received a low platform solubility score (0.23) and integrated score (39.5). The surface-arginine-optimized variant SrMilN_SASAR showed clearly improved solubility experimentally relative to the parent - enough that a reasonable amount of protein entered the soluble fraction - and was scored higher by the platform (solubility 0.57; integrated score 55.7). Importantly, this should not be overstated as a purification or production success: the soluble fraction still presented a family of bands, and the material was not usable as a clean, single-species reagent for downstream work. Thus the model correctly ranked the engineered, partially solubilized variant above the insoluble parent, consistent with the experimental redesign, while the wet-lab outcome remains an incomplete rescue. Figure 2B places this single-pair comparison in the broader external-test context by showing the confusion matrix at threshold 0.46 (TP = 13, FP = 2, TN = 13, FN = 10), clarifying that directional ranking on SrMilN/SASAR coexists with remaining false negatives in the mid-score range. Full gel images and quantitative band densitometry will be required for peer-reviewed reporting of this case.

### Open statement of dataset limits

Primary quantitative calibration used eSOL cell-free E. coli solubility and yield rather than intact-cell industrial fermentation (Figure 1A). Most public training sequences lack coding DNA, leaving codon- and rhythm-dependent modules weakly supervised. The collaborator external set underlying Table 2 and Figure 1B-C is small (n = 38), project-enriched, and yields wide confidence intervals; sensitivity (56.5%) is only modestly above chance. Binary soluble/insoluble labels omit quantitative yield and catalytic activity. Offline EC-class labeling is at chance (3/20 = 15%; Table 1) and is not a manufacturability claim or marketed capability. Public industrial ranking used literature priors rather than matched assay measurements under identical conditions, and the Spearman interval on n = 10 includes zero. Future DNA-linked expansion sources include construct-level expression panels such as public kinase E. coli expression libraries and NESG/SoluProt resources after license confirmation.

As summarized in Table 1, the group ordering provides preliminary directional evidence that the integrated score can track some relative expression difficulty, but the Spearman confidence interval is wide and includes no association. These results do not establish calibrated prediction of soluble titre or activity because the categories were qualitative literature-derived priors rather than measurements obtained under standardized experimental conditions. The heuristic EC classifier agreed with the curated UniProt/BRENDA first-level class for only 3 of 20 enzymes (15%; 95% CI 5.2–36%), at or below six-class chance; EC output is therefore demoted to an optional provisional label and is not presented as a platform capability.

### Case study: Bacillus licheniformis α-amylase

Bacillus licheniformis α-amylase (UniProt P06278; curated EC 3.2.1.1) was selected as a familiar starch-processing enzyme representing the tractable literature category. The unified workflow produced a baseline score of 61.87 (good) and an integrated score of 46.30 (fair). RR-motif and ribosome-stall risks were low, whereas the solubility module returned 0.0 (poor). The provisional EC classifier assigned class 2 rather than curated class 3. Thus, the overall score retained a broadly favorable relative position within the rank subset, but the enzyme-level output also exposed two model weaknesses: non-discriminatory solubility scoring and inaccurate EC assignment.

### Case study: Taq DNA polymerase

Thermostable DNA polymerase I from Thermus aquaticus (UniProt P19821; curated EC 2.7.7.7) was selected as a contrasting thermophilic enzyme representing the condition-dependent literature category. It received a baseline score of 39.50 (poor) and an integrated score of 24.30 (poor). Eight RR motifs and high predicted stall risk contributed to the low integrated assessment, while the provisional EC class correctly matched class 2. The result demonstrates how the platform can flag sequence-level translational risks even for an enzyme that is producible industrially under optimized conditions. It should not be interpreted as evidence that Taq polymerase cannot be expressed in E. coli.

### Observed limitations of the retrospective benchmark

The public ranking benchmark (Table 1) remains useful only for relative ordering hypotheses and has clear limits. Literature tractability categories were not controlled for vector, strain, induction temperature, fusion tag, secretion route, cofactor conditions, or assay. Protein-only public inputs incompletely exercise codon harmonization and translation-rhythm modules. The first-level EC heuristic is at chance (15%) and is retained solely as a non-capability disclosure. Earlier panel runs that forced solubility to 0.0 for every protein are superseded by the calibrated solubility model in Figure 1A and by the external holdout in Table 2 and Figures 1B–C and 2, which themselves remain small-n estimates with wide intervals. Independent host-matched validation using measured soluble yield or catalytic activity, DNA-linked constructs, and larger multi-project cohorts remains required before the integrated score can be interpreted as a quantitative predictor of fermenter titre.

## Discussion

### Industrial enzyme development: from discovery to manufacturability

Industrial enzyme programs operate under constraints that differ fundamentally from academic protein expression studies. A candidate enzyme may exhibit excellent specific activity, thermal stability, and substrate selectivity under screening conditions yet fail to progress through manufacturing gates because recombinant titre, soluble yield, or batch-to-batch consistency does not meet commercial thresholds (Schmid et al., 2001; Bornscheuer et al., 2012). The global industrial enzyme market exceeded USD 7 billion in recent estimates, with hydrolases, oxidoreductases, and transferases accounting for the majority of commercial products (Adrio and Demain, 2014; Panke and Wubbolts, 2005). In detergent, food processing, and pharmaceutical intermediate manufacturing, cost per kilogram of active enzyme directly determines process economics (Link et al., 2013; Turner and O’Reilly, 2013); expression failure at the construct design stage therefore carries disproportionate financial risk relative to the cost of the computational screen that might have prevented it.

### The gap between codon optimization vendors and bioprocess reality

Commercial gene synthesis workflows typically offer codon optimization as a default service, often maximizing CAI or eliminating rare codons without regard to translational rhythm, co-translational folding kinetics, or the phylogenetic distance between native producer and expression host (Gustafsson et al., 2004; Cannarozzi et al., 2015; Plotkin and Kudla, 2011). Empirical studies have shown that synonymous codon changes can alter protein structure and function, challenging the assumption that codon choice affects only translation rate (Agashe et al., 2013; Zhou et al., 2013). For industrial enzymes sourced from actinomycetes and filamentous fungi - which together supply a large fraction of hydrolases, oxidoreductases, and transferases used in biocatalysis (Ferrer et al., 2016; Adrio and Demain, 2014) - this one-size-fits-all approach frequently produces constructs that amplify rather than resolve expression problems. The representative S. hygroscopicus α-amylase case (UniProt P08486; GenBank M15540) illustrates this disconnect: baseline metrics indicated a fair expression band (59.1) with low RR burden but weak E. coli codon adaptation and high GC (70%), while integrated co-translational analysis revealed folding bottlenecks and solubility-patch constraints that would likely manifest as inclusion bodies or low soluble titre in a 10–50 L pilot fermenter (Rosano and Ceccarelli, 2014; de Groot and Goverde, 2012). An industrial team relying solely on CAI-based gene design would have insufficient visibility into these failure modes prior to purchase order placement.

### Strategic value across the industrial enzyme value chain

From an industry perspective, the platform addresses three high-value decision points in the enzyme development pipeline (Nielsen et al., 2011; Woodley, 2013):

- Lead prioritization: Metagenomic and genome-mining campaigns routinely yield dozens to hundreds of candidate sequences (Ferrer et al., 2016; Steele et al., 2009). Computational pre-screening by expression manufacturability, rather than catalytic potential alone allows R&D teams to rank constructs before committing to gene synthesis, cloning, and shake-flask expression.
- Host selection and construct design: Relocating a Streptomyces hydrolase to E. coli versus Pichia pastoris versus B. subtilis involves different codon profiles, secretion pathways, and glycosylation constraints (Rosano and Ceccarelli, 2014; Baeshen et al., 2015; Branduardi et al., 2004). Parameterized host and source organism contexts enable side-by-side comparison of expression risk before host commitment (Demain and Vaishnav, 2009).
- Technology transfer and scale-up readiness: When an enzyme program transitions from research to process development, regulatory and quality teams require documented rationale for construct design (Woodley, 2013). Structured platform output component diagnostics, harmonization recommendations, and variant proposals provides traceable decision support that complements experimental expression data in tech transfer packages.

### Industry-specific enzyme classes and expression challenges

Different industrial enzyme categories present distinct manufacturability profiles. Technical-grade hydrolases (amylases, proteases, lipases) used in detergents and food processing are typically required at high volumetric productivity (Adrio and Demain, 2014); even moderate solubility penalties can translate to economically unacceptable downstream processing costs (Kyte and Doolittle, 1982; Magnan et al., 2009). Pharmaceutical and fine-chemical biocatalysts often oxidoreductases or transferases requiring cofactors, metal centers, or disulfide bonds face additional constraints captured by EC class context and co-translational folding analysis (Bornscheuer et al., 2012; Sheldon and Brady, 2022). Enzymes from extremophile sources frequently carry GC-rich coding sequences and arginine-rich surface patches that stall ribosomes when transferred to mesophilic production hosts (Anne et al., 2014; Atomi et al., 2011). Local analysis without mandatory external database calls is particularly relevant for companies protecting proprietary sequence data during early-stage evaluation.

### Integration into industrial workflows

We envision the platform as a gate in the standard industrial enzyme workflow: sequence acquisition → computational manufacturability assessment → ranked variant design → gene synthesis order → micro-scale expression validation → fermenter scale-up (Terpe, 2006; Rosano and Ceccarelli, 2014). Secure local execution supports evaluation of sequences that cannot be uploaded to cloud-based annotation services. The platform does not replace experimental expression testing, fermentation optimization, or downstream formulation development (Demain and Vaishnav, 2009; Garcia-Fruitos, 2012). Rather, it reduces expensive experimental iterations by front-loading sequence-level risk assessment at the point where redesign is cheapest, consistent with design–build–test–learn paradigms in bioprocess development (Nielsen et al., 2011).

### Limitations and future directions from an industry standpoint

Industrial adopters should recognize current limitations. Sequence-centric scoring does not incorporate host-strain background, promoter strength, induction temperature, or fusion-tag strategy, all of which significantly affect industrial titre (Rosano and Ceccarelli, 2014; Terpe, 2006). Optional EC-class labels and organism predictions require curated-database confirmation; EC heuristics are not a validated capability (UniProt Consortium, 2023). Current evidence, as quantified in Tables 1–3 and visualized in Figures 1–2, supports only preliminary screening utility on small cohorts: held-out eSOL Spearman correlation of approximately 0.55 (Figure 1A); external wet-lab balanced accuracy 71.6% (57.4–84.4%) and MCC 0.43 (0.15–0.69) (Table 2; Figure 1B), with specificity stronger than sensitivity (56.5%, near chance on this set) and directional calibration at the score extremes (Figure 1C). The SrMilN→SASAR pair (Table 3; Figure 2A) illustrates directional ranking after a partial solubility rescue (multiple soluble bands; not purification-ready), while Figure 2B shows that false negatives persist. Near-term scientific priorities include expansion with DNA-linked construct panels, replacement or removal of the EC heuristic in favor of curated annotation lookup, tighter coupling to fermentation titre models, and continued external testing while permanently excluding collaborator holdout data from model fitting.

As a longer-term industrial future-work plan, we aim to expose the analysis modules as Model Context Protocol (MCP) tools so that laboratory or enterprise agentic AI systems can invoke manufacturability scoring, case-study retrieval, UniProt-linked sequence ingest, and variant comparison inside controlled design-build-test-learn loops without uploading proprietary enzyme sequences to third-party model APIs. Concretely, planned MCP tools include expression analysis, case-study listing and detail retrieval, accession-based sequence fetch, side-by-side variant rescoring, and condensed explanations of limiting score components. Building on that interface, we plan a thin agent harness - not unconstrained autonomous biology - that (i) ingests a candidate CDS or accession, (ii) calls MCP analysis tools, (iii) proposes a small set of surface-arginine and rhythm-preserving harmonization edits grounded in module recommendations, (iv) re-scores the redesigned constructs, and (v) stops when a performance band improves or a maximum iteration budget is reached, emitting an auditable pre-synthesis report. Two deployment modes are envisioned for partners: bring-your-own-agent, in which customer agents (for example Cursor-, Claude-, or LangGraph-based planners) call our local MCP server behind the firewall; and a hosted, tool-constrained agent that may call only approved MCP endpoints with authentication, credit limits, and run history suitable for technology-transfer packages. In all cases the agent remains human-in-the-loop decision support for construct triage; it does not replace experimental expression testing or fermenter commitment.

## Conclusion

Recombinant expression remains the principal bottleneck between industrial enzyme discovery and commercial biocatalytic deployment (Schmid et al., 2001; Bornscheuer et al., 2012; Adrio and Demain, 2014). Conventional codon optimization, while necessary, is insufficient to predict manufacturability of enzymes relocated from phylogenetically distant native producers to standard industrial expression platforms (Gustafsson et al., 2004; Cannarozzi et al., 2015; Plotkin and Kudla, 2011). We present an integrated computational platform that combines baseline sequence metrics with co-translational modeling to provide structured, interpretable expression risk assessment for industrial enzyme candidates.

The platform accepts diverse sequence inputs, operates locally without external database dependencies, and returns baseline and integrated composite assessments, component-level diagnostics, harmonization recommendations, variant design proposals, and visualization outputs within a single unified analysis workflow. Representative analysis of the authentic S. hygroscopicus α-amylase CDS (UniProt P08486; GenBank M15540) demonstrates how integrated co-translational assessment reveals manufacturability constraints invisible to CAI-based assessment alone (Welch et al., 2009; O’Brien et al., 2014). Preliminary public ranking (Table 1), eSOL calibration (Figure 1A), and a small independent wet-lab external holdout with confidence-bounded sensitivity, specificity, MCC, and calibration (Table 2; Figure 1B–C), including the SrMilN arginine-surface directional partial-rescue example (Table 3; Figure 2), support transparent partner-facing evaluation. Offline EC labeling is disclosed as at-chance and is not claimed as a capability. Openly stated dataset limits clarify that the current evidence base is decision-support grade rather than a guarantee of fermenter titre.

From an industry perspective, the platform addresses a well-documented gap between sequence acquisition and fermenter commitment: the need for rapid, secure, and economically rational construct prioritization in enzyme programs where expression failure carries disproportionate cost (Panke and Wubbolts, 2005; Link et al., 2013; Woodley, 2013). By embedding co-translational biology into a practical pre-synthesis review workflow (Zhang et al., 2009; Netzer and Hartl, 2016), the platform supports more informed decision-making across lead prioritization, host selection, and technology transfer. Experimental validation remains essential (Demain and Vaishnav, 2009); this tool is designed to ensure that validation effort is directed toward the most promising constructs. Proprietary scoring parameters and implementation details are intentionally withheld in this preprint to protect intellectual property while communicating scientific scope, industrial utility, and representative performance. In the retrospective public-data benchmark, integrated scores followed the expected tractable > condition-dependent > difficult group ordering and showed a moderate rank association with literature-derived difficulty (ρ = 0.458), while important limitations in EC classification and solubility discrimination were identified. These findings justify further independent experimental validation but do not establish calibrated prediction of expression yield.

